# Chemical defense and tonic immobility in early life stages of the Harlequin cabbage bug, *Murgantia histrionica* (Heteroptera: Pentatomidae)

**DOI:** 10.1101/2021.01.29.428818

**Authors:** Eric Guerra-Grenier, Rui Liu, John T. Arnason, Thomas N. Sherratt

## Abstract

Antipredation strategies are important for the survival and fitness of animals, especially in more vulnerable life stages. In insects, eggs and early juvenile stages are often either immobile or unable to rapidly flee and hide when facing predators. Understanding what alternative antipredation strategies they use, but also how those change over development time, is required to fully appreciate how species have adapted to biotic threats. *Murgantia histrionica* is a stink bug, conspicuously colored from egg to adult, known to sequester defensive glucosinolates from its cruciferous hosts as adults. We sought to assess whether this chemical defense is also present in its eggs and early nymphal instars and quantified how it fluctuates among life stages. In parallel, we looked at an alternative antipredation strategy, described for the first time in this species: tonic immobility. Our results show that the eggs are significantly more chemically defended than the first two mobile life stages, but not than the third instar. Tonic immobility is also favored by hatchlings, but less so by subsequent instars. We argue the case that over development time, tonic immobility is a useful defensive strategy until adequate chemical protection is achieved over an extended feeding period.

## Introduction

Animals are faced with constant threats, such as predation and parasitism, and have to overcome them in order to survive and reproduce. Several antipredation strategies have evolved in numerous taxa in order to minimize predation pressures. One of them is the use of chemical compounds to deter predators through their unpalatability and/or toxicity. Species using such chemical defenses frequently advertise their unprofitability through warning signals (e.g. conspicuous body coloration), allowing predators to learn to avoid defended prey when exposed to the signals through associative learning: a phenomenon called aposematism (Poulton 1890; Skelhorn et al. 2016). Although usually studied in active life stages, chemical defenses in insects are also known to occur in eggs (Guerra-Grenier 2019). In their thorough book chapter on the subject, Blum and Hilker (2002) distinguish between two types of chemical protection in insect eggs: the defensive compounds can either be produced autogenously by the parents (intrinsic origin) or can be sequestered from their diet (extrinsic origin), although some species are known to do both, such as *Photuris* fireflies (González et al. 1999). An example of eggs protected by *de novo* chemicals is found in the Australian chrysomelid *Paropsis atomaria* Olivier 1807 (Nahrstedt and Davis 1986; Blum and Hilker 2002). All life stages of these beetles possess cyanogenic glygosides which, when under attack, can be modified to release hydrogen cyanide (HCN), a known respiratory inhibitor. As their *Eucalyptus* host plants are free of cyanogenic glycosides, it is believed that the chemical defense is produced by the insects themselves and then transferred into the eggs. Eggs protected through sequestration can be found for example in two orthopteran species of the genus *Poecilocerus* feeding on milkweed plants: *P. bufonius* (Klug 1832) and *P. pictus* (F. 1775) (Fishelson et al. 1967; Pugalenthi and Livingstone 1995; Blum and Hilker 2002). Milkweeds are famous for their poisonous cardenolides, which are sequestered by the aposematic *Poecilocerus* and later incorporated in their eggs. Furthermore, whether they are of intrinsic or extrinsic origin, chemical defenses can sometimes be provided to the eggs by the fathers (Eisner et al. 2002). This is potentially advantageous to both parents since females must only pay a fraction of the metabolic costs for synthesis and/or sequestration while males with lots of toxins may be favored during mate choice.

Aside from chemical protection, tonic immobility (TI) is another common antipredation strategy. Individuals engaging in tonic immobility enter a completely motionless state after physical contact with potential predators. TI is also referred to thanatosis or death feigning in the literature, because immobile prey are reminiscent of deceased individuals (Ruxton 2006). However, as pointed out by Humphreys and Ruxton (2018*a*, and references therein) in their recent review (and noted by Darwin (1883)), death feigning individuals often adopt postures that are dissimilar to those of dead individuals of the same species. Henceforth, we will use the term *tonic immobility.* TI has been hypothesized to be an efficient anti-predation strategy through various mechanisms (not necessarily mutually exclusive) including mimicking death, the signaling of chemical defenses (Miyatake et al. 2004, 2009; Ruxton 2006) and the reduction of attack rates by motion-oriented predators (Edmunds 1974; Prohammer and Wade 1981; Miyatake 2001). Although it is well known, tonic immobility is still relatively understudied and its use is most likely under-reported (Humphreys and Ruxton 2018*a*).

A good candidate for the study of antipredation strategies is the Harlequin cabbage bug, *Murgantia histrionica* (Hahn 1834) (Heteroptera: Pentatomidae). This species of stink bug feeds mainly on cruciferous plants (Brassicaceae), but is known to have over 50 possible host plants (McPherson 1982). Native to Mexico, it is established in several U.S. states and is occasionally observed in Canada (McPherson 1982; Paiero et al. 2013). It is a recognized pest of economically important crops such as cabbage, collard, kale and broccoli (Ludwig and Kok 2001; Wallingford et al. 2011) and is thus studied for its management, more recently in a semiochemical context (Conti et al. 2003; Peri et al. 2016). Part of what explains the success of these bugs is their low number of natural enemies; they have a few parasitoids and virtually no predators or competitors (McPherson 1982; Amarasekare 2000), suggesting efficient anti-predation strategies. Contrary to most North American pentatomids, which are cryptic brown or green, *Murgantia histrionica* is easily recognizable through its orange, white and black markings (Figure 1a). Aliabadi et al. (2002) suggested that this conspicuous color pattern acts as a warning signal to reduce predation by birds. Indeed, they showed that, much like some other species feeding on cruciferous plants such as aphids and sawflies (Opitz and Müller 2009), adult Harlequin bugs can sequester glucosinolates from their host plant into their body tissues. Glucosinolates (mustard oil glycosides) are secondary metabolites produced by cruciferous plants and their hydrolysis products (isothiocyanates, nitriles, etc.) are toxic and used as antiherbivory defenses (Louda and Mole 1991; Opitz and Müller 2009). Although only described in *Murgantia histrionica*, this chemical defense could potentially be present in other members of the Strachini tribe (i.e. other *Murgantia spp., Eurydema spp., Stenozygum spp*., etc.) given their shared body coloration (orange, white and black markings) and use of glucosinolate-producing host plants (i.e. Brassicaceae, Capparaceae, etc.) (Exnerová et al. 2008; Samra et al. 2015). While it is still unknown whether the stink bugs sequester, along with glucosinolates, the enzyme myrosinase responsible for the hydrolysis of the compounds, or if they produce an enzyme *de novo*, a chemical reaction during the act of predation is necessary for mustard oils to be effective against predators (Aliabadi et al. 2002).

**Figure 1.**
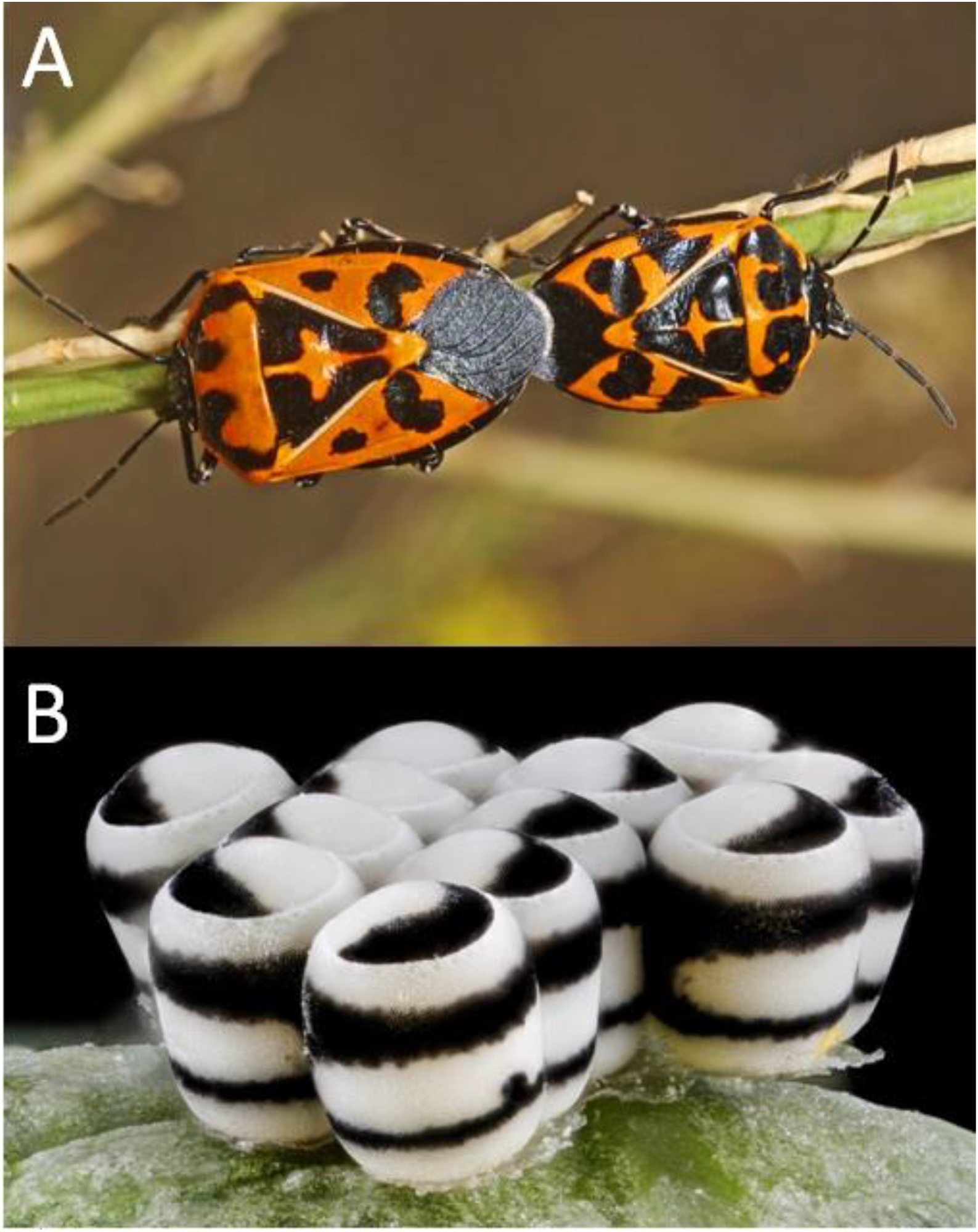
A) Adults (photo by Judy Gallagher (CC BY 2.0)) and B) eggs (Public Domain Mark 1.0) of *Murgantia histrionica*.

Much like the adults of the species, *M. histrionica* eggs and nymphs are also quite distinctive. The eggs (Figure 1b), usually laid in clutches of 12 (two rows of six eggs), are white with two black bands and a black spot on the external side. The operculum is also black and white, and the black portion can either form a ring or be shaped as to resemble the Yin-Yang symbol. Like in the adults, the egg color pattern is generally conserved throughout the Strachini tribe (Kalender 1999; Samra et al. 2015). Nymphal coloration is somewhat similar to that of the adults in terms of hues and pattern complexity, but white markings are also found on their dorsal side. Although sequestration of glucosinolates presumably starts during immature stages, it is currently unknown whether nymphs and eggs are actually chemically defended against vertebrate and invertebrate predators. Furthermore, tonic immobility has been observed in hatchlings that were physically disturbed (Guerra-Grenier, personal observations). This is interesting because, like in other stink bug species, this life stage does not feed (Canerday 1965; Zahn et al. 2008), meaning that if glucosinolates are not vertically obtained through parental bestowal, chemical defense cannot be achieved until later on in their development. TI could thus be an alternative line of defense, possibly used to deter motion-oriented predators.

The aim of the present study was to assess the extent of antipredation defenses in eggs and early nymphal instars of the Harlequin cabbage bug. First, we asked if the white portion of the egg color pattern, already thought to be conspicuous against a green crucifer leaf background, allows for an increased contrast (i.e. more intense warning signal) by also reflecting wavelengths in the ultraviolet (UV) part of the electromagnetic spectrum. Indeed, several insect species can perceive UV light and incorporate ultraviolet coloration in their signals (Cronin and Bok 2016) and leaves are known to absorb ultraviolet radiation from the sun (Gutschick 1999). Secondly, we wanted to assess whether eggs and nymphs have glucosinolates and, if so, whether their concentrations differ among life stages. We predicted that concentrations in nymphs increase over successive molts, reflecting an increased sequestration over feeding time. Finally, we wanted to investigate the use of TI in nymphs by looking at its frequency and duration, predicting that it is used by the insects in order to compensate for lower chemical defenses and that TI use decreases as glucosinolate sequestration increases.

## Materials and Methods

### Insect colony

*M. histrionica* individuals were collected in the Beltsville area (Maryland, USA) in 2016 and subsequently cultured at 28 ± 1°C and a 16L: 8D light cycle in an incubator (VWR Chamber Diurna Growth 115V, Cornelius, USA) supplied with a jar of water for added moisture. Adults were reared continuously in 30 cm^3^ ventilated polyester cages (BugDorm, Taichung, Taiwan) while nymphal instars were kept in ventilated plastic containers (length: 22 cm; width: 15 cm; height: 5 cm; Rubbermaid, USA). All were fed with rinsed fresh store-bought broccoli *(Brassica oleracea* var. *italica*), replaced three times per week. Eggs were collected three times per week from the sides of the cages and broccoli and put into Petri dishes (diameter: 9cm) lined with a filter paper until hatching, at which point they were transferred to the plastic containers.

### Ultraviolet photography

Ten clutches (totaling 106 eggs), one adult male and one adult female were photographed under both ultraviolet and visible light using a Nikon D70 camera with an El-Nikkor 80mm lens and a Baader U UV filter (Baader Planetarium, Germany: 310-390nm UV transmission). The El-Nikkor lens is sensitive to wavelengths 320nm and higher (Verhoeven and Schmitt 2010). A MTD70 EYE color arc bulb (70W, 1.0A power source, www.eyelighting.co.uk) was used as the only light source and was chosen for its D65 spectrum (the same spectrum as sunlight). Eggs were glued on double sided tape on a Petri dish (diameter: 9cm) and pictures were taken from above and from the external (banded) side. Adults were photographed on their ventral and dorsal sides.

### Phytochemical analyses

A total of 301 individual Harlequin cabbage bugs were sampled from the lab colony for glucosinolate extractions, following an adapted version of the protocol established by Doheny-Adams et al. (2017). Individuals were first separated into life stages (whole eggs, first instars, second instars and third instars), subsequently split into triplicates of equivalent mass (3 x ~18.00 mg for eggs; 3 x ~13.00 mg for first instars; 3 x ~40.00 mg for second and third instars) and macerated in a solution of 70% methanol at a dilution ratio of 1 ml of solution per 4 mg of insect mass. The shells left behind by the first instars, which were collected before they started feeding, were also pooled (~3.00 mg per replicate) and treated the same way. Flowerets of broccoli (440.45 mg) bought from the same batch the insects fed on were ground with liquid nitrogen and then transferred into 70% methanol at the same dilution ratio mentioned above. Once prepared, samples were put into a water bath at 70°C for 30 minutes for extraction.

The resulting extracts were filtered through a 0.22 mm PTFE filter and analyzed on a Shimadzu UPLC-MS system (Mandel scientific company Inc, Guelph, Canada) which contains LC30AD pumps, a CTO20A column oven, a SIL-30AC autosampler and a LCMS-2020 mass spectrometer. Briefly, 1 μl of each fraction was injected through the autosampler to a Luna omega polar C18 column (100 x 2.1mm, 1.6 μm particle size, Phenomenex, Torrance, USA). Mobile phases were H2O and acetonitrile, with 0.1% formic acid in both. The isocratic elution method was initialed with 2% acetonitrile for 5 minutes. The column was then washed with 100% acetonitrile for 3 min and re-equilibrated for 5 minutes before the next injection. The flow rate was set at 0.5 ml/min with the column tempered at 5Ü°C.

The mass spectrometer with electrospray ionization (ESI) interface operated in negative selective ion monitoring (SIM) mode, the nebulizing gas flow was set at 1.5 L/min and drying gas flow was at 10 L/min. The desolvation line temperature and heat block temperature were set at 250°C and 400°C respectively. The detector was monitoring m/z at 436 [M-H]- with 938 u/sec scan speed. Linear calibration curves were built by injecting dilutions of glucoraphanin (Cayman chemicals, Ann Arbor, USA) (Figure 2) which bracketed the compound concentration in the samples. Calibration curves were prepared at five concentration levels (0.2-10 ng on column) and R^2^ values obtained.

**Figure 2.**
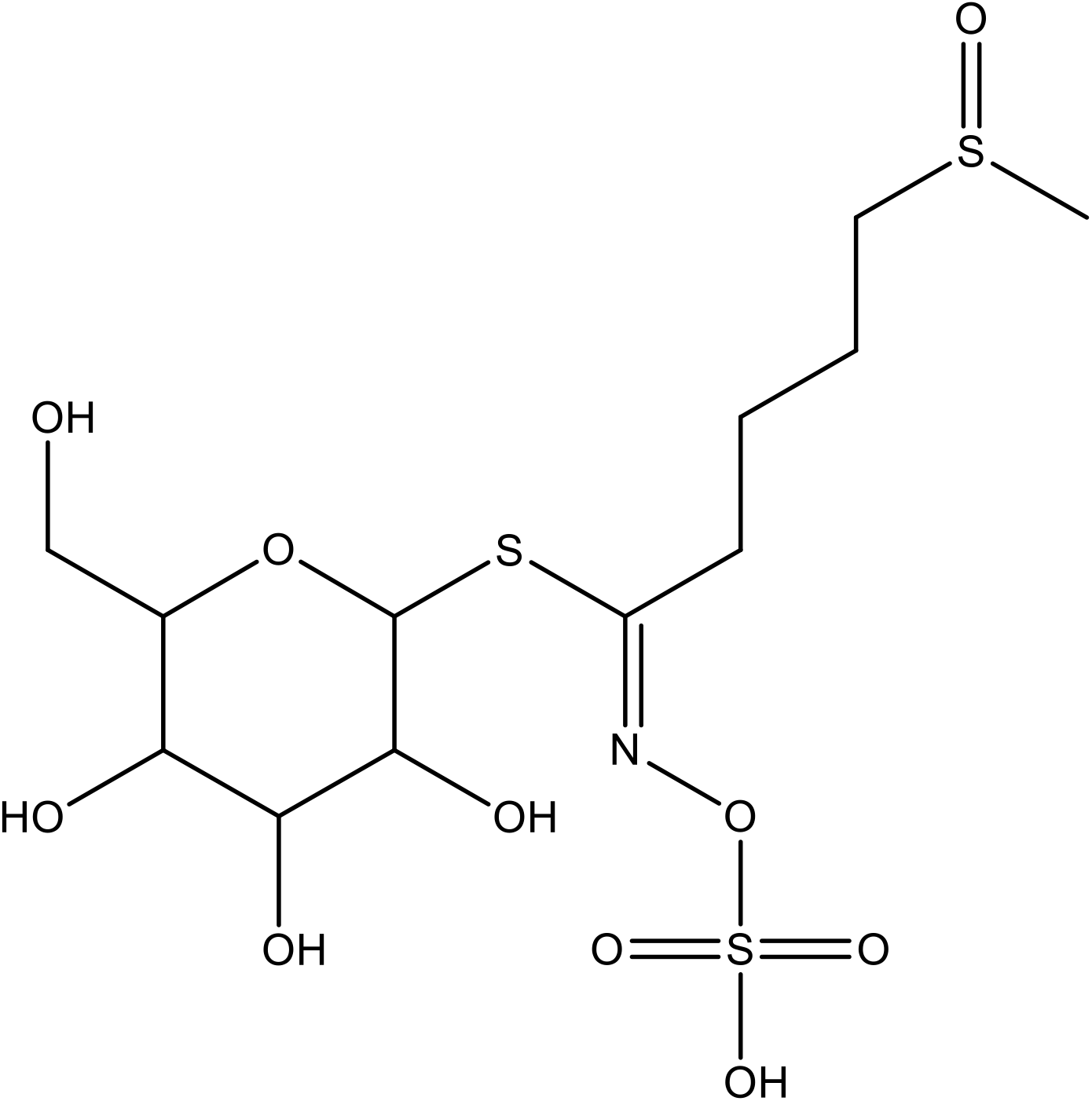
Schematic representation of a glucoraphanin molecule.

### Behavioral assays

Eggs were sampled from the lab colony and put in clutch-specific Petri dishes (diameter = 9 cm). Individuals that hatched from those eggs were reared in the same environmental condition as the general rearing until they reached the first (86 from 8 clutches), second (81 from 10 clutches) and third instars (97 from 14 clutches). The number of clutches used increased with increasing instar as to obtain similar sample sizes while compensating for the baseline mortality rate. One at a time, every individual went through the following protocol. Bugs were first placed on a filter paper (diameter: 12.5 cm). They were then prodded by one of us (EG-G) using a paint brush. Each trial consisted of up to five sessions of five prods (25 in total), with each session separated by one minute. The presence of TI and its duration (if present) was recorded. Trials ended when individuals either displayed TI postures (Figure 3) for 5 seconds or more, or if all 25 prods unsuccessfully elicited the behavior. TI postures kept for less than 5 seconds were not considered given that such a short duration could be due to sensory overload or to a period of physiological recovery from impact. The clutch of origin of every individual, as well as the order at which they were taken from their clutch-specific container and prodded, were also recorded. A Petri dish lid of the same size as the filter paper was placed onto the arena between prodding sessions in order to stop the bugs from escaping. The soft hair of the brush was frequently moistened, twisted together and then dried by contact with another filter paper in order to reduce the risk of trapping the bugs among the brush hair. All trials were carried out by the same observer.

**Figure 3.**
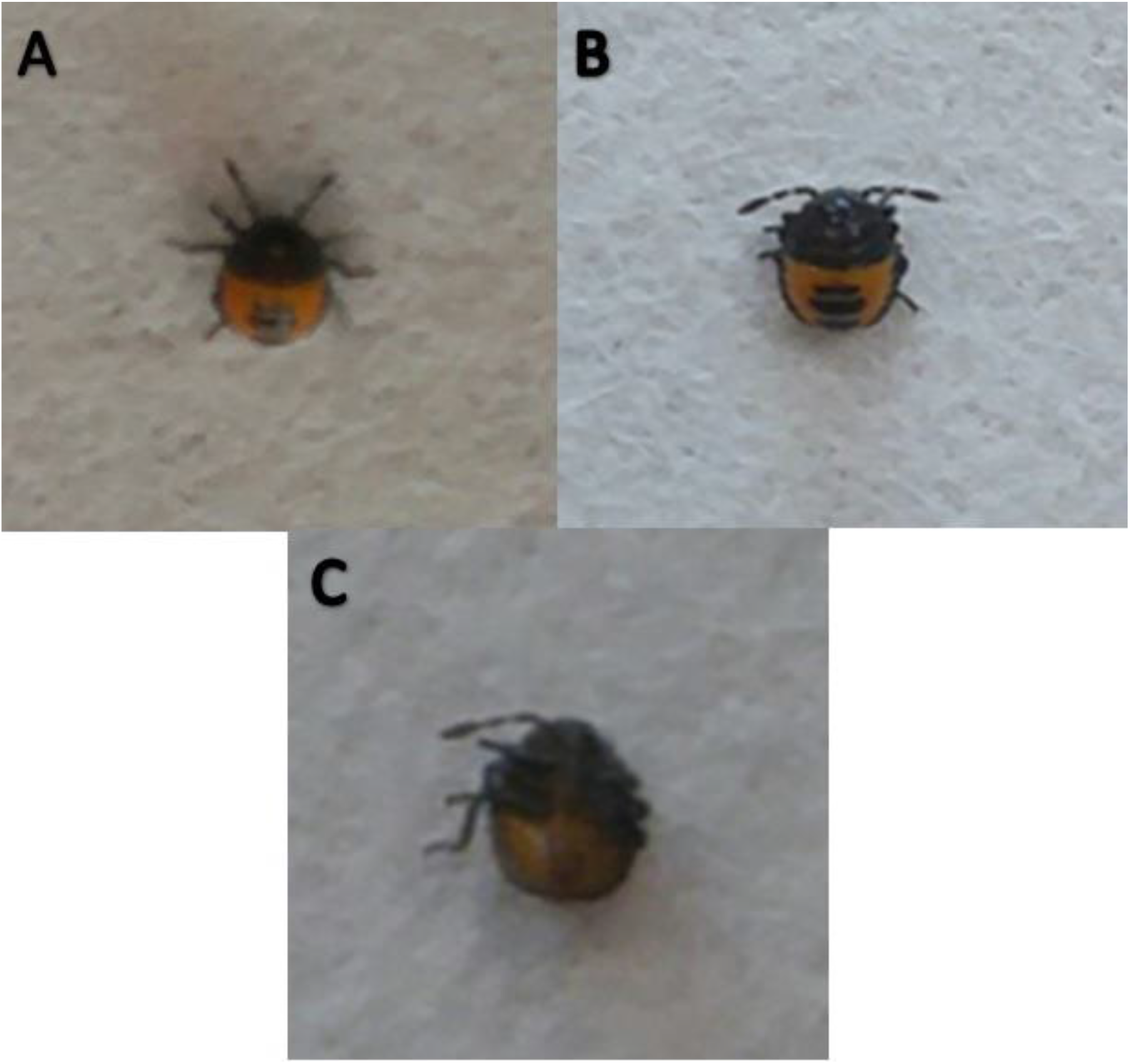
Postures adopted by *M. histrionica* nymphs during behavioral assays. Nymphs could be active (A) or display tonic immobility when on their ventral (B) or dorsal (C) side. The pictures shown here are of an individual in its first instar. Photo credits: Eric Guerra-Grenier.

### Statistical analyses

All analyses were conducted with R version 3.3.3 (R Core Team 2017). A Kruskal-Wallis rank sum test allowed for the comparisons of glucoraphanin concentrations among eggs and the three nymphal instars. Dunn’s tests were subsequently used for *post-hoc* pairwise comparisons between life stages using the “dunn.test” package in R (Dinno 2017). Differences in concentrations between hatchlings and their eggshells were analyzed by fitting a linear model following a square root transformation of the concentration data to successfully ensure homogeneity and normality.

Generalized linear mixed models (GLMMs) and generalized linear models (GLMs) with binomial error distributions were used to regress the probability of tonic immobility as a function of instar and prodding order (fixed factors), with clutch of origin as a random factor. These models were fitted using the “lme4” package in R (Bates et al. 2014). General linear mixed models (LMERs) and linear models (LMs) were fitted to test for the effect of the same factors on the duration of tonic immobility, assuming normally distributed error. For all statistical models that we fitted to the TI data, the significance of each of the fixed factors was determined with likelihood ratio tests using the “car” package in R (Fox and Weisberg 2011). AIC values finally determined which models among those based on all or a subset of the factors best fitted the data (Table 1). Simultaneous tests for general linear hypotheses with Tukey contrasts were subsequently used for post-hoc pairwise comparisons between nymphal instars using the “multcomp” package in R (Hothorn et al. 2008).

**Table 1.**
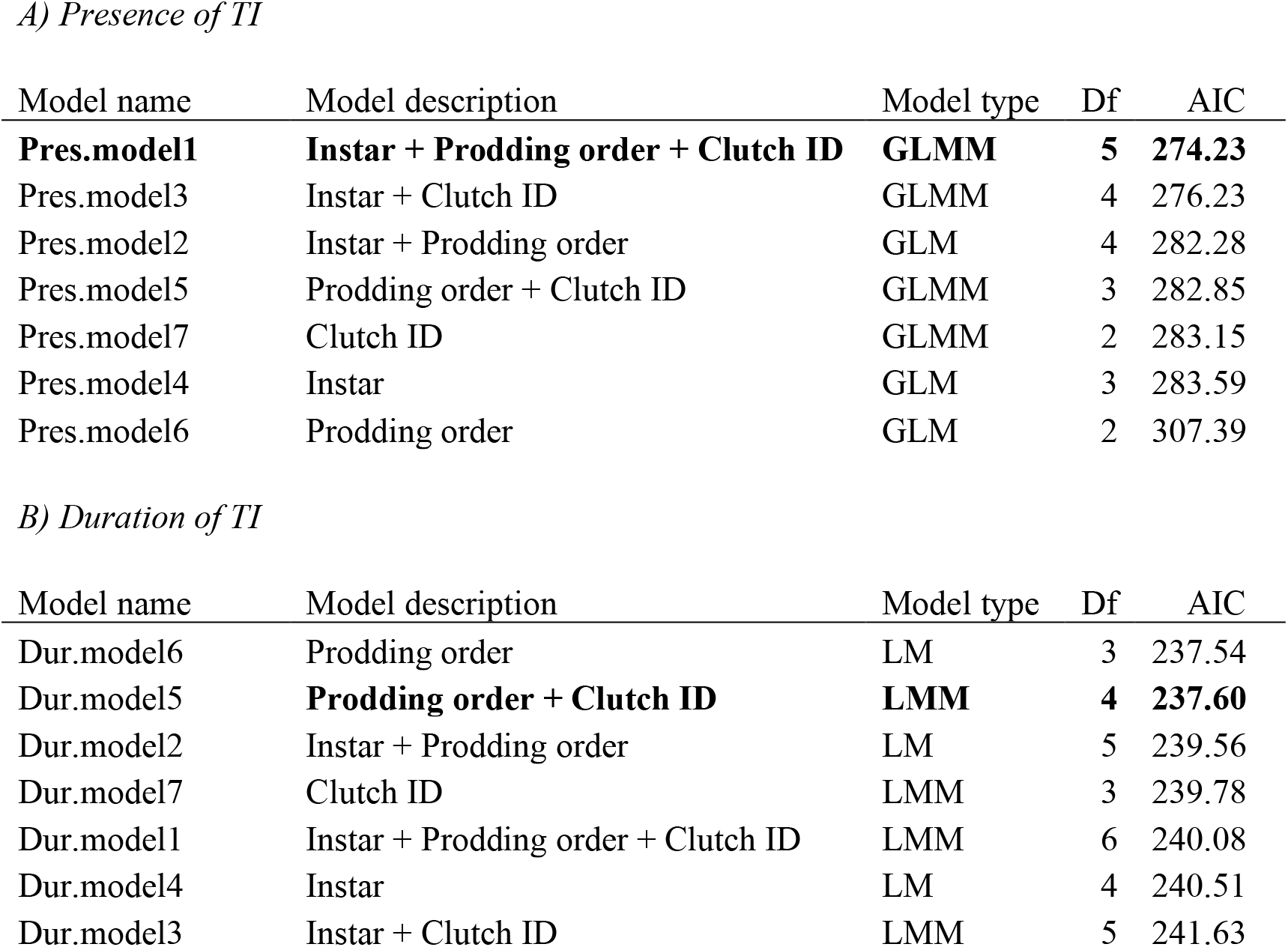
Comparison of the fit of the models testing for the effects of fixed and random factors on A) the presence and B) duration of TI displayed by nymphs. Rows in bold indicate which models best fitted the data based on AIC values. If two or more models shared the lowest AIC (± 0.1), the one with the most degrees of freedom was chosen.

| Model name | Model description | Model type | Df | AIC |
| --- | --- | --- | --- | --- |
| **Pres.model1** | **Instar + Prodding order + Clutch ID** | **GLMM** | **5** | **274.23** |
| Pres.model3 | Instar + Clutch ID | GLMM | 4 | 276.23 |
| Pres.model2 | Instar + Prodding order | GLM | 4 | 282.28 |
| Pres.model5 | Prodding order + Clutch ID | GLMM | 3 | 282.85 |
| Pres.model7 | Clutch ID | GLMM | 2 | 283.15 |
| Pres.model4 | Instar | GLM | 3 | 283.59 |
| Pres.model6 | Prodding order | GLM | 2 | 307.39 |

*B) Duration of TI*
| Model name | Model description | Model type | Df | AIC |
| --- | --- | --- | --- | --- |
| Dur.model6 | Prodding order | LM | 3 | 237.54 |
| Dur.model5 | **Prodding order + Clutch ID** | **LMM** | **4** | **237.60** |
| Dur.model2 | Instar + Prodding order | LM | 5 | 239.56 |
| Dur.model7 | Clutch ID | LMM | 3 | 239.78 |
| Dur.model1 | Instar + Prodding order + Clutch ID | LMM | 6 | 240.08 |
| Dur.model4 | Instar | LM | 4 | 240.51 |
| Dur.model3 | Instar + Clutch ID | LMM | 5 | 241.63 |

## Results

### UV reflectance in eggs and adults

Eggs exposed to ultraviolet light (Figure 4b, d) showed high reflectivity in the pale, but not dark markings of the color pattern in all ten clutches photographed. The same level of contrast was present under visible light only (Figure 4a, c). This implies that the pale markings of the eggs are perceived as true white to UV-sensitive animals as well as to those who are not (e.g. humans). Unlike the eggs, both the male and the female adults showed almost no reflection of wavelengths between 320-390nm (Figure 4e-l). This suggests that the features (i.e. pigments and/or refracting nanostructures) responsible for the white coloration perceived by humans in adults and eggs are different to some extent.

**Figure 4.**
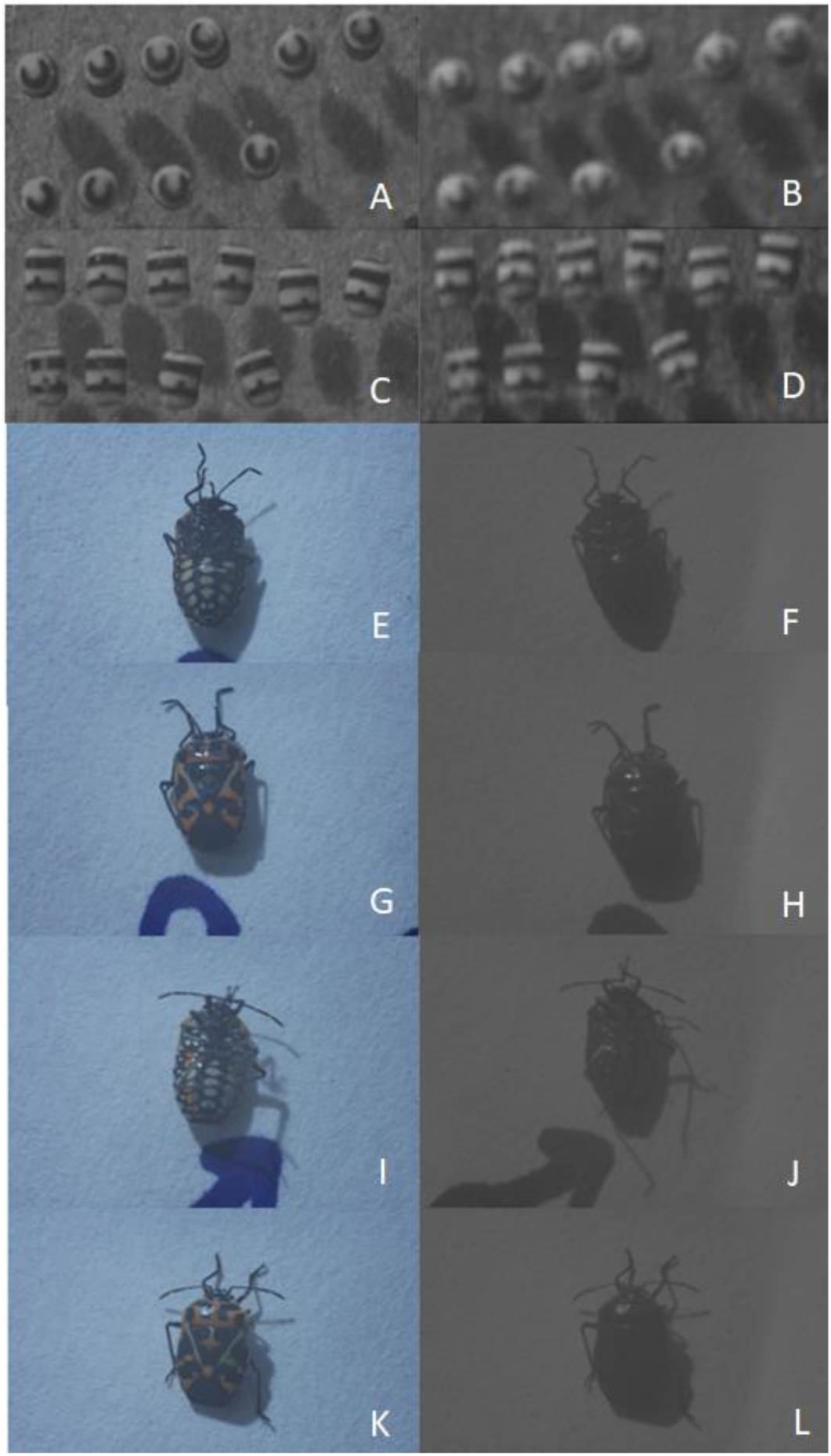
Pictures of eggs and adults taken in the visual (left panels) and ultraviolet (right panels) spectra. Eggs are seen from their top (A-B) and external sides (C-D). The female is seen from her ventral (E-F) and dorsal (G-H) sides. The male is seen from his ventral (I-J) and dorsal (K-L) sides. UV pictures were converted in black and white images so that UV light could be perceived by human eyes on a luminance scale (the brighter the markings, the more UV-reflecting they are). Photo credits: Eric Guerra-Grenier.

### Chemical defenses in eggs and nymphs

Glucoraphanin was detected in broccoli as expected, but also in the eggs and hatchlings (first instars) (Figure 5). Simple qualitative (presence/absence) detection of the compound is likely to be misleading for second and third instars, as its presence in the gut from previous feeding events make it impossible to know if glucoraphanin is also present in body tissues. Concentrations of glucoraphanin were subsequently evaluated in eggs and all three instars (Figure 6a) and were shown to differ among life stages (H = 8.7436, df = 3, p = 0.0329). Post-hoc comparisons revealed that eggs had significantly higher concentrations than first (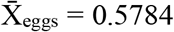, S_eggs_ = 0.0833, 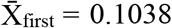, S_first_ = 0.1011, Z = 2.6042, p = 0.0046) and second instars (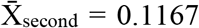, S_second_ = 0.0073, Z = 2.3778, p = 0.0087), but did not differ from third instars (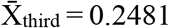, S_third_ = 0.5294, Z = 1.323, p = 0.1288). Yet, third instars had similar levels of glucoraphanin to first (Z = −1.4720, p = 0.0755) and second instars (Z = −1.2455, p = 0.1065). Concentrations in the first two nymphal instars did not differ significantly (Z = −0.2265, p = 0.4104). Since hatchlings had significantly less compound than eggs, their concentrations were compared to those of the shells they left behind when hatching (Figure 6b). The analysis revealed that, of all the compound present in the eggs, most of it is laced within the shells (LM: F = 88.002, df = 1, 4, p < 0.001), leaving only a small portion of glucoraphanin available for integration by the developing insects.

**Figure 5.**
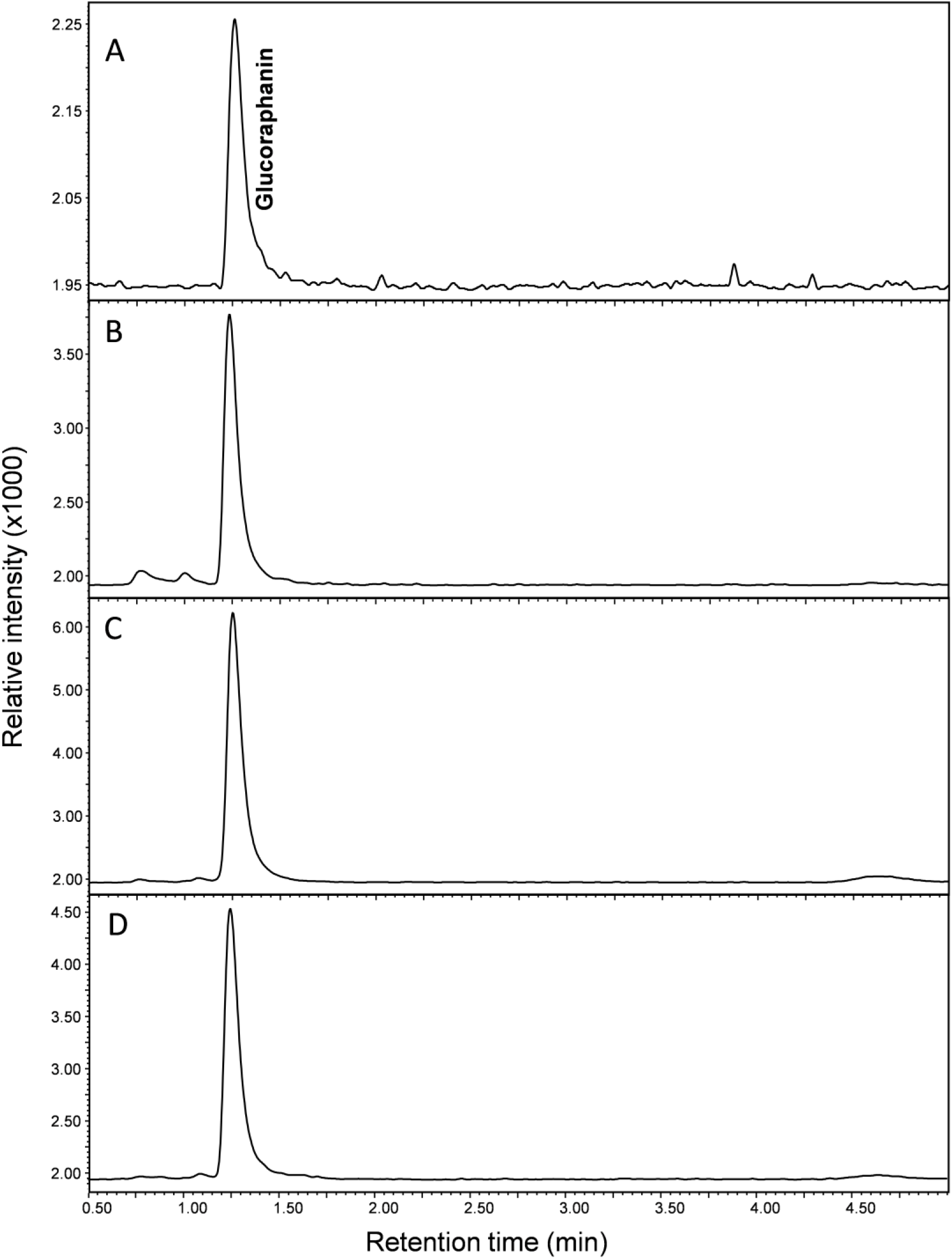
UPLC-ESI/MS negative extracted ion chromatogram of glucoraphanin (m/z = 436 (M-H)-). A) Glucoraphanin commercial standard; B) Broccoli flowerets; C) Whole eggs; D) First instars (hatchlings).

**Figure 6.**
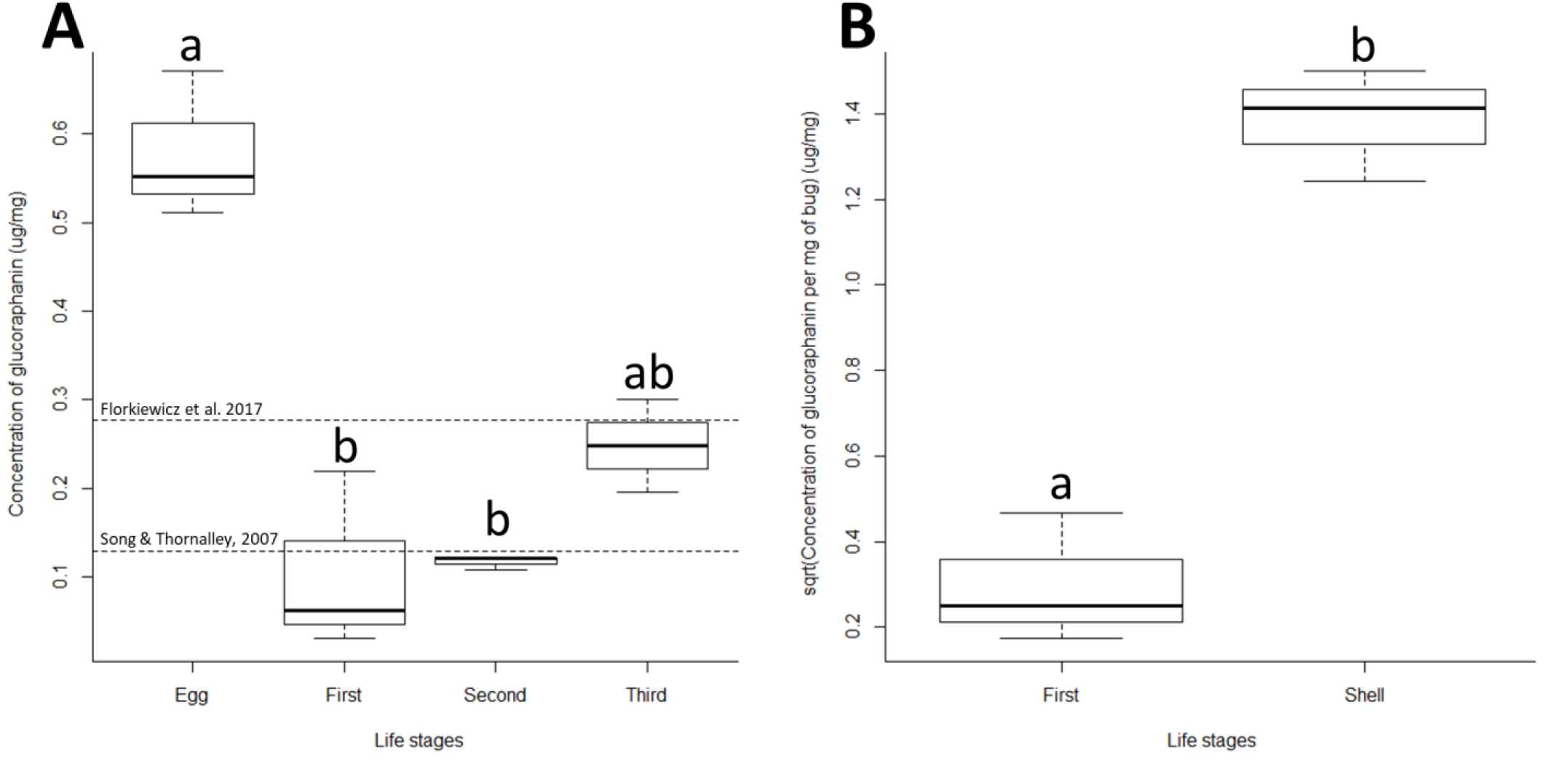
Boxplot representations of glucoraphanin concentrations in various life stages. A) The concentrations of glucoraphanin (μg/mg) in whole eggs, first, second and third nymphal instars chemical defenses. The dotted lines represent concentrations of glucoraphanin in broccoli (reported by Florkiewicz et al. (2017) and Song and Thornalley (2007)) as a means of comparison. B) The concentrations of glucoraphanin (μg/mg) in hatchlings (first instars) and their shells.

### Tonic immobility in nymphs

The probability of displaying tonic immobility of a given nymph was strongly influenced by which instar it was in (GLMM: χ^2^ = 13.0001, df = 2, p = 0.0015): hatchlings displayed the behavior more frequently than second (p = 0.0304) and third instars (p = 0.0013), whereas the later two instars had similar display frequencies (p = 0.5120) (Figure 7a). The probability of displaying TI also reduced the later nymphs were sampled from their clutch of origin (GLMM: χ^2^ = 3.8653, df = 1, p = 0.0493) (Figure 7b). The duration of the behavior was shorter in individuals tested later within a clutch compared to those tested earlier (LMM: χ^2^ = 4.2635, df = 1, p = 0.0390), but was not affected by nymphal instar. The clutch from which nymphs originated was also a key factor partially predicting both the probability of displaying TI (Table 2A) and the duration of the behavior (Table 2B).

**Figure 7.**
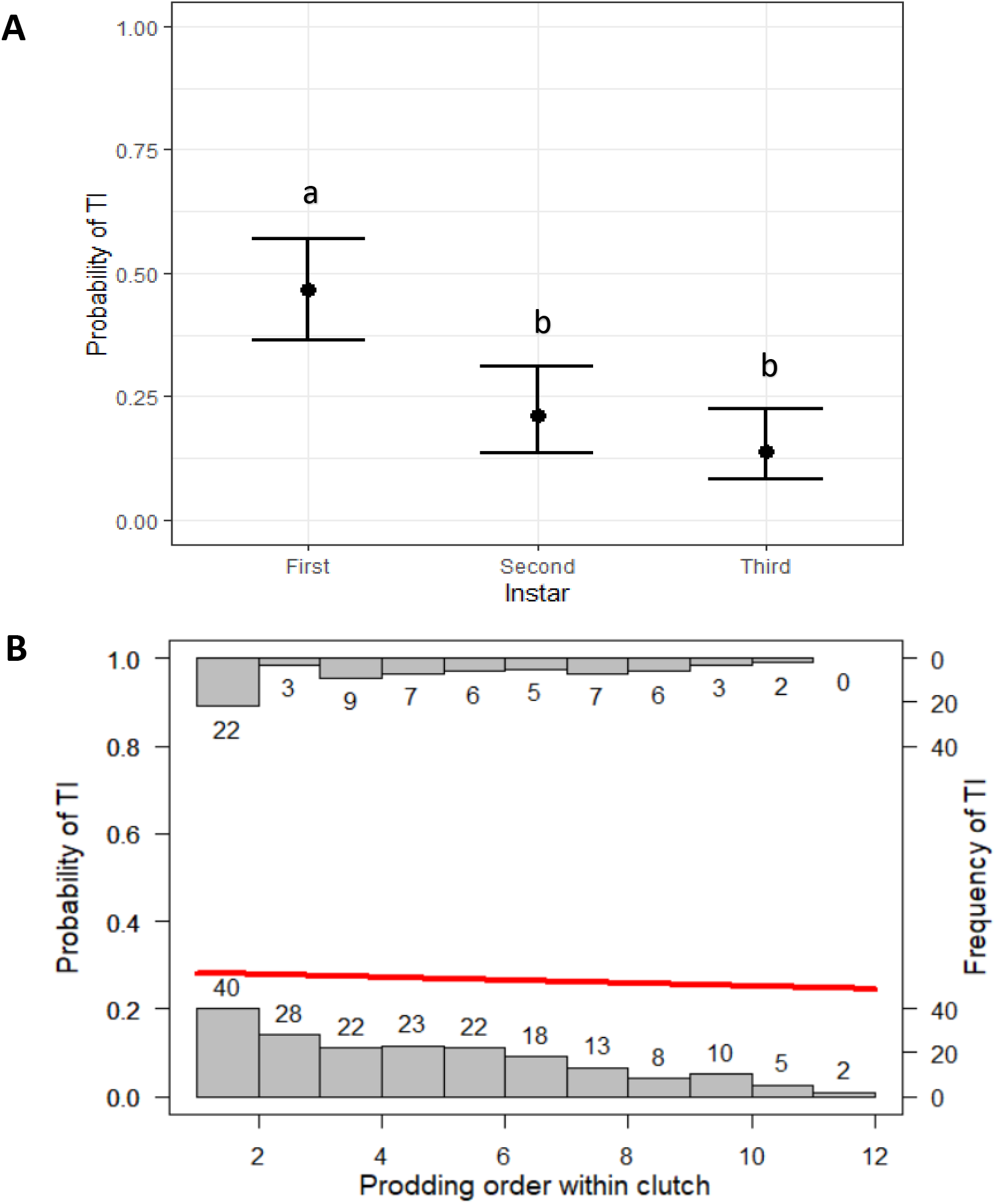
A) The average probability of entering tonic immobility across instars. Error bars represent 95% binomial confidence intervals while letters represent differences in statistical significance. B) The effect of prodding order within a clutch on an individual’s probability of entering tonic immobility across all instars. Histograms represent the frequencies of TI while the red trendline shows variation in the probability of entering TI as a function of prodding order.

## Discussion

The aim of this study was to investigate two potential antipredation strategies in eggs and juvenile *Murgantia histrionica*: chemical protection and tonic immobility. Our results suggest that both strategies are used by the bugs, but also that their use varies across life stages. Given that eggs are completely immobile and are “sitting ducks” to would-be predators, behavioral strategies are unavailable, leaving only chemical protection as a viable defense. This could explain why concentrations in the eggs are much higher than in the young nymphs, in which glucoraphanin levels do not vary significantly among instars. However, levels of compound in the eggs are similar to those of third instars. Given that first and second instars have a significantly lower concentration than eggs, we can speculate that third instars are at an intermediate level, indicating that sequestration probably starts at this stage but has not yet reached the degree seen in adults. Indeed, according to Aliabadi et al. (2002), adults sequester glucosinolates in their tissues at concentrations 20-30 times higher than those found in the gut (i.e. about the same concentration as in cruciferous plants). Our results however demonstrate that concentrations in nymphs are similar to concentrations in broccoli (Song and Thornalley 2007; Florkiewicz et al. 2017, although variation occurs among cultivars as shown by Brown et al. 2002). This may be because it takes time to build up the concentration in body tissues. It may also be because small nymphs of true bugs, including the Harlequin cabbage bug, are exposed to different predators than adults and bigger nymphs (mostly invertebrate vs. mostly vertebrate respectively) (Exnerová et al. 2003), and these predators likely have different susceptibilities to mustard oils. Although both vertebrates and invertebrates are usually intolerant to glucosinolates (Halkier and Gershenzon 2006), such compounds are used by plants to limit insect feeding (Louda and Mole 1991) while they can potentially be beneficial to vertebrates at low enough concentrations (Shapiro et al. 1998).

Interestingly, a large proportion of the glucosinolates in the eggs is laced within the shells and is thus not accessible to the bugs for integration before hatching. This could explain why, even though egg predators have seldom been reported, eggs are often attacked by parasitoid wasps of the genera *Trissolcus* (Platigastridae) and *Ooencyrtus* (Encyrtidae) (McPherson 1982; Amarasekare 2000; Conti et al. 2003; Peri et al. 2016). These endoparasitoids drill through the eggshell with their ovipositor and lay their eggs directly within the host eggs, meaning that they bypass most of the chemical defense. Furthermore, even though the larvae enter in contact with the lower dose of glucosinolates within the eggs, parasitoids are known to be more specialized on their hosts than predators are on their prey and may have evolved an increased resistance to the compounds. An alternative hypothesis to explain the higher concentration of compound in the shells could be that mothers use them as toxin sinks. By doing so, females would reduce their metabolic investment in detoxification while still avoiding compromising the development of their offspring. Eggshells sequestration as a detoxification mechanism has previously been suggested in bird eggs (Kitowski et al. 2014).

Unlike for chemical protection, Harlequin bug nymphs showed life stage-specific variation in their probability, but not duration, of entering tonic immobility. Indeed, first instars displayed the behavior significantly more frequently than second and third instars. As we have not been able to show that instar-related variation in glucoraphanin content occurs, we cannot confirm our hypothesis that TI is present in younger life stages to compensate for the lack of chemical defense. A higher probability of TI in hatchlings could however be explained by variation in life history traits. *M. histrionic*a is not really active until its second instar. Zahn et al. (2008) reported that neonate bugs do not feed and remain aggregated with their siblings; only after the first molt did they start to feed and disperse. In other words, if attacked, individuals in their first instars are too small and probably not active enough to successfully flee. TI thus becomes the optimal behavioral strategy, in addition to chemical protection. It is also conserved at a lower degree in second and third instars. Yet again, variation in life history traits could explain the lower frequency in older nymphs: fleeing becomes a safer option as size and speed increase, leading up to an almost zero probability of entering TI in the final nymphal instars and adults (Guerra-Grenier, personal observations).

Regardless of instar, TI use was also influenced by the order in which we prodded individuals of a given clutch. Those tested first had a higher probability of entering tonic immobility than those tested last, and kept the posture for a longer period. This may be because of an alarm cue perceived by the nymphs. Such a cue could be the visual detection of conspecifics being “attacked”, but is most likely olfactory in nature. There is a reason why stink bugs are named this way: they release pungent volatile chemicals when disturbed, and these chemicals are known to elicit dispersal in neighboring conspecifics (Ishiwatari 1974). In *Murgantia histrionica* specifically, some of these compounds are thiocyanides derived from the glucosinolates they absorb from their host plants (Aldrich et al. 1996; Aliabadi et al. 2002). It would make sense for individuals reacting to these chemical cues to engage less frequently and in shorter TI periods if they are stimulated to flee. It should however be noted that ecological factors cannot fully explain the use of TI. Inter-clutch variation in the probability of displaying TI and its duration suggests that there are other factors affecting tonic immobility in *M. histrionica*. One possibility is that the propensity to engage in TI is partly hereditary, which would not surprising considering that genetic variation in TI is known to occur in other taxa (Miyatake et al. 2004). However, we cannot confirm this hypothesis just yet given that nymphs used in this experiment were collected from different areas in the colony (e.g. plant stem vs cage walls) and at different times, possibly adding unaccounted for environmental variation in the mix.

As mentioned in the introduction, TI is sometimes positively correlated with chemical defenses and has been hypothesized to act as a warning display (Miyatake et al. 2004, 2009; Ruxton 2006). Such a correlation is not apparent in *Murgantia histrionica* based on our results: the frequency of TI diminishes from first to second and third instar, but glucosinolate levels remain the same across these three life stages. With that said, given that adults have much higher concentrations compared to what we observed in early nymphs (Aliabadi et al. 2002) and that they do not seem to use TI as an antipredation strategy (Guerra-Grenier, personal observation), there is probably a negative correlation between chemical defense and tonic immobility across all life stages, from hatchlings to adults. In this case, TI would not serve as a warning signal but rather simply as an alternative defense strategy, potentially deterring motion-oriented predators. More data on TI and glucosinolate content in fourth and fifth instar as well as in adults need to be collected in order to test for the existence of such a relationship.

In addition to the description of their chemical defense, we have demonstrated that *M. histrionica* eggs are most likely highly conspicuous to predators, whether they are UV-sensitive or not. Indeed, the white markings of the eggs reflect ultraviolet light, meaning that they appear white (i.e. they reflect all the wavelengths that a given visual system is sensitive to) to any potential predator, regardless of whether they are sensitive to wavelengths between 300-400nm. Such a color pattern thus contrasts strongly against a leaf background, which usually absorbs UV-light (Gutschick 1999). In a general sense, it is safe to assume that *M. histrionica* eggs are aposematic in that they are both conspicuous to would-be predators, and unprofitable; as such, the black and UV-white pattern probably advertises glucosinolate-related unpalatability or toxicity. However, further experiments measuring prey (eggs, but also nymphs) acceptance by ecologically relevant predators over repeated exposures are necessary to confirm that predators indeed learn to avoid consumption of the bugs based on recognition of their color pattern. Such tests should also compare predation levels by chewing vs. piercing-sucking predators to test for the relative importance of the shell (which composes most of the chemical defense) in the learning process. Confirmation of efficacy is also required for the TI strategy: although the posture adopted by Harlequin bug nymphs when disturbed fits with the defining characteristics of TI (Humphreys and Ruxton 2018*a*), it is still unknown whether it reduces predation success in this species. An alternative explanation for the behavior could be that it would allow the bugs to fall off their host plant when threaten in nature, a defensive strategy described in other insect taxa (Gross 1993; Chaboo 2011; Humphreys and Ruxton 2018*b*). Further testing of TI in *M. histrionica* thus requires using bugs placed on their host plants in order to confirm whether immobilized individuals fall off the plant or remain close to predators on the leaves.

Complex color patterns often play a role in multiple defensive and/or communication strategies (Marshall 2000; Tullberg et al. 2005; Caro et al. 2016, 2017). Specifically for striped black and white patterns, known functions vary from crypsis (e.g. disruptive camouflage, countershading, etc.) to warning signals and can interfere with a predator’s ability to attack a prey through a dazzling effect (Stevens et al. 2008; Izzo et al. 2014; Feltwell 2016). Aside from aposematism, the complex *M. histrionica* egg color pattern may thus have alternative adaptive functions, and future studies should look into those in this species as well as in taxa with similar egg coloration such as *Eurydema* spp. and *Stenozygum spp.* (plesiomorphy: Kalender 1999; Samra et al. 2015), but also in *Piezodorus spp* (convergent evolution: Bundy and McPherson 2000). For example, the stripes and dot may convey an intraspecific signal influencing the ovipositional behaviors of conspecific females, provide disruptive camouflage through distance-dependent crypsis, protect against harmful solar radiation and/or reduce desiccation (Guerra-Grenier 2019). Additionally, because eggs vary in their proportion of black, pigment variation may help with heat absorption in colder periods. Such a thermoregulatory function has actually been shown in the mobile life stages of the species, where late nymphal instars exposed to colder temperatures will metamorphose into adults with a higher degree of melanisation than those exposed to warmer temperatures (White and Olson 2014).

## Acknowledgements

We would like to thank Paul K. Abram, Sophie Potter, Changku Kang, Felipe Dargent, Casey Peet-Paré, Tammy Duong, Yolanda Yip, Ian Dewan, Lauren Efford, Karl Loeffler-Henry, Gustavo L. Rezende, Greg Bulté, Naomi Cappuccino, Mark Forbes and Andrew Simons for helpful discussions and/or technical assistance, as well as Donald C. Weber for sending us eggs of *Murgantia histrionica* to build our lab colony.

## Declarations

### Author contributions

Conceptualization, EG-G, RL, JTA, TNS; Methodology, EG-G, RL, JTA, TNS; Investigation, EG-G, RL; Formal Analysis, EG-G, RL, TNS; Writing – Original Draft, EG-G, RL; Writing – Review & Editing, EG-G, RL, JTA, TNS; Supervision, JTA, TNS; Funding Acquisition, JTA, TNS.

